# Adaptive spatiotemporal filtering of social electrosensory signals by cerebellar feedback

**DOI:** 10.64898/2026.09.28.755101

**Authors:** DR Miller, EK Shinn, B Harrison, G Marsat

## Abstract

Sensory feedback can suppress predictable input, yet most research on this mechanism used simplified stimuli that do not capture the spatial and temporal structure of natural signals. We examined how cerebellum-like feedback, in brown ghost knifefish, shapes the encoding of realistic social electrosensory signals. These naturalistic signals replicated the spatial structure of social signals, spanned the range of typical AM beat frequencies, and incorporated movement-related contrast envelopes. Single-unit recordings were obtained from electrosensory lateral line lobe pyramidal cells while a conspecific-like signal was delivered. Responses were compared before and after pharmacological blockade of descending cerebellar feedback. As expected, feedback strongly attenuated responses to spatially-realistic low-frequency beats but did not alter beat-cycle coding at higher frequencies. However, feedback reduced the neural representation of movement-related contrast envelopes across all beat frequencies. Using information-theoretic tools, we showed that feedback strength adapts dynamically to the stimulus contrast over the past few hundred ms. Furthermore, we examined how the spatial coding by the pyramidal cell population is influenced by feedback. We show that feedback enhances localization cues. This improvement arose because feedback reduces weak background responses while not affecting the strongest responses, thereby increasing spatial contrast. These findings show that the same descending pathway jointly filters temporal and spatial features of the full range of natural signals, adapting to recent input statistics. This process exemplifies a general mechanism by which a predictive feedback can improve the efficiency of natural sensory representations.

## Introduction

Natural sensory signals are structured across space and time. Rather than encountering isolated changes in stimulus intensity, animals typically experience complex scenes composed of spatial gradients, local contrast, temporal fluctuations, and movement-generated dynamics. A major challenge for sensory circuits is therefore to preserve behaviorally relevant variation while reducing responses to stimulus components that are redundant, predictable, or weakly informative. This principle has been studied in various sensory systems (Smith and Lewicki 2006; Uchida et al. 2014). In vision, for example, the statistics of natural scenes have been used to explain center-surround receptive fields, spatiotemporal filtering, and the decorrelation of retinal ganglion cell responses (Karamanlis et al. 2022; Pitkow and Meister 2012). In some sensory systems, much of what is known about sensory filtering has come from simplified stimuli, such as full-field flashes, gratings, or spatially uniform modulations. Although processing of naturalistic signals has been investigated before, we still have an incomplete understanding of how filtering mechanisms identified under controlled conditions operate when behaviorally relevant signals vary simultaneously across multiple dimensions.

Feedback pathways are well positioned to implement this type of adaptive filtering because they can combine incoming sensory activity with information about stimulus context, recent history, and behavioral state. In predictive coding frameworks, feedback signals are often proposed to carry expectations that suppress predictable components of sensory activity while preserving responses to deviations from those expectations (Bastos et al. 2012; Keller and Mrsic-Flogel 2018). The comparison of a prediction with an actual signal is a generic computation central to the cerebellar and cerebellar-like circuits, where granule cell–parallel fiber pathways involves diverse contextual signals that can be used to generate predictions of expected sensory input (Bell et al. 2008). These circuits therefore provide experimentally tractable systems for asking how feedback reshapes sensory representations.

The electrosensory system of weakly electric fish has been especially useful for studying this problem because the relevant sensory signals, feedback pathways, and neural responses can be fully characterized. Wave-type gymnotiform fish continuously generate an electric organ discharge (EOD) and detect perturbations of this self-generated field through electroreceptors distributed across the skin. These afferents project topographically to pyramidal neurons in the electrosensory lateral line lobe (ELL), whose apical dendrites receive extensive descending input from cerebellum-like granule cell–parallel fiber pathways (Bell and Maler 2005; Krahe and Maler 2014). Previous work has shown that this feedback can generate negative images of predictable electrosensory input, thereby reducing responses to redundant modulations and improving the representation of behaviorally relevant signals (e.g. Bastian 1986; Bol et al. 2011; Hofmann and Chacron 2019). This extensive body of work has characterized this feedback circuit at the level of identified cells, plasticity rules, and population responses, providing an unusually detailed mechanistic framework for asking how the same feedback operates when processing natural conspecific signals with complex spatiotemporal structures.

Conspecific signals in wave-type gymnotiforms provide a natural context in which sensory input varies across several behaviorally relevant dimensions at once. When two fish are near one another, the interference between their electric organ discharges produces amplitude modulations, or beats, whose frequency is determined by the difference between the two discharge frequencies (Zakon et al. 2002). These beat signals are not static in overall strength (the peak-to-trough contrast of the beat). As fish move relative to one another, changes in distance and orientation alter the strength of the beat and generate slower envelope modulations that can carry information about relative motion and distance (Fotowat et al. 2013; Stamper et al. 2013; Yu et al. 2012). In addition, the electric field generated by a nearby conspecific is not spatially uniform across the focal fish’s body surface (in this paper, the term “focal fish” refers to the individual for which we are considering the sensory process). Instead, it creates an electric image whose amplitude varies across the receiver’s body and the receptor array, producing spatial gradients that can support localization and shape population activity patterns across the ELL (Milam et al. 2019; Ramachandra et al. 2026). Thus, natural conspecific signals combine beat frequency, envelope dynamics, and spatial structure, making them well suited for testing how cerebellar feedback filters sensory input with realistic temporal and spatial structures.

Most studies of cerebellar feedback in the gymnotiform ELL have focused on how the circuit attenuates low-frequency, spatially diffuse modulations. This work established that feedback can reduce responses to redundant beat signals and identified cellular and synaptic mechanisms that constrain this cancellation to relatively slow temporal modulations (Bol et al. 2011). This framework leaves two related questions unresolved. First, many natural social interactions generate high-frequency beats whose individual cycles are too fast to be canceled, but whose amplitude envelopes vary on slower timescales that are encoded by ELL pyramidal neurons and may still engage feedback (Fotowat et al. 2013; Middleton et al. 2006). Second, feedback effects have often been characterized using spatially uniform stimuli that drive the feedback pathway strongly. Indeed, this feedback pathway is driven by topographically broad inputs and therefore does not operate strongly for spatially localized stimuli such as a small prey (Berman and Maler 1999; Hofmann and Chacron 2019). Spatially realistic conspecific signals produce electric images with strong spatial gradients across the receptor array and are thereby neither small localized signals nor global uniform signals (Ramachandra et al. 2026). It therefore remains unclear how strongly feedback affects the responses to conspecific signals and whether it shapes the pattern of responses across the population to preserve, or even enhance, the spatial information carried by these signals.

Within a predictive-feedback framework, we could expect this pathway to act on both spatial and temporal aspects of natural signals to improve the efficiency with which they are represented. Here, we determine how cerebellar feedback shapes the encoding of realistic conspecific signals with naturalistic spatial and temporal characteristics. We recorded ELL pyramidal cell responses in *Apteronotus leptorhynchus* while presenting realistic conspecific signals that have spatially realistic structures, a wide range of beat frequencies as observed naturally, and modulated contrast envelopes as experienced by interacting fish. By comparing responses with feedback intact and after pharmacological blockade of descending input, we characterize its impact on the coding of realistic conspecific signals. We show how the feedback adapts to recent stimulus history to shape the responses to envelope modulation for both low- and high-frequency beats. Furthermore, we show that it affects most strongly the weaker responses within the population, thereby enhancing spatial contrast and localization cues. Our results highlight a general principle in sensory processing whereby adaptive filtering of sensory signals improves the efficiency with which natural signals are encoded.

## Material and Methods

### Animals

Experiments were performed in 38 wild-caught brown ghost knifefish (*Apteronotus leptorhynchus*) obtained from commercial suppliers. Fish of various sizes were housed in small groups in 30-gallon aquaria maintained at 26–28°C and a water conductivity of 200–400 μS/cm. The sex and age of the animals were not determined. All procedures were approved by the West Virginia University Institutional Animal Care and Use Committee and followed institutional guidelines for animal care and use.

### Surgical preparation

Surgical procedures followed established methods for in vivo recordings from the electrosensory lateral line lobe (Allen and Marsat 2019; Bastian 1986; Bol et al. 2011). Fish were anesthetized with tricaine methanesulfonate (Syndel) delivered in oxygenated water passed continuously over the gills. Once surgical anesthesia was established, the skin overlying the posterior portion of the skull was removed, and a local anesthetic was applied to the exposed tissue. The skull was secured to a rigid post, and a small craniotomy was made to provide access to the ELL. Fish were immobilized by injection of ∼1 µg/g gallamine triethiodide (Sigma-Aldrich).

Following surgery, anesthesia was discontinued by switching the respiratory flow to anesthetic-free water and local anestetic was applied regularly to the incised skin. The fish was positioned in a 45 × 45 × 20 cm experimental tank, with the body submerged, with only the cranial opening at the top of the head maintained above the water surface. The head remained fixed throughout the experiment, and oxygenated water was continuously passed over the gills. Recordings were therefore obtained from an awake, immobilized preparation. Tank water was maintained at approximately 27 ± 1°C and at a conductivity of 300 μS/cm.

### Electrophysiological recordings

Single-unit extracellular recordings were obtained using custom electrodes (Carlson and Kawasaki 2004; Dowben and Rose 1953; Heiligenberg and Rose 1986). Sharp glass micropipettes were made with a Flaming/Brown P-1000 puller (Sutter Instrument). The tip was broken manually to a diameter of ∼5 µm. The tip was then filled with melted Woods Metal (a bismuth-based alloy) and connected to a metal wire. The tip was electroplated with gold and then platinum black, giving the tip a large surface for a small volume, thus maximizing the spatial and electrical properties of the electrode. Electrode signals were amplified with a Model 1700 differential amplifier (A-M Systems) and digitized at 20 kHz using a Digidata 1500 data-acquisition interface and AxoScope software (Molecular Devices). Recordings were stored for subsequent spike detection and offline analysis.

Recordings targeted ON-type pyramidal cells in the lateral and centrolateral segments of the ELL (LS and CLS). These segments are strongly involved in processing conspecific signals (Marsat et al. 2009; Metzner and Juranek 1997). Within each segment, we focused on superficial and intermediate pyramidal cells because their apical dendrites receive parallel-fiber input from the eminentia granularis posterior (EGp) and are therefore influenced by the indirect cerebellar feedback pathway examined here (Bastian et al. 2004). The stereotyped surface vasculature and the anatomical relationship of the ELL to the overlying cerebellum guided electrode penetrations. Segment and laminar position were estimated from the mediolateral and rostrocaudal location of the penetration and the depth of the electrode below the dorsal brain surface (Maler et al. 1991). These assignments were also supported by response properties measured with step, sinusoidal, and random amplitude modulations. We compared their temporal tuning, spontaneous firing rate, and phasic or phasic–tonic response profile to published characteristics (Krahe et al. 2008).

The classical receptive field of a subset of neurons was localized to ensure that the sampled cells covered the whole rostro-caudal and ventral-dorsal range of the fish’s body ipsilateral to the recording side. To do so, stimuli were delivered using a small dipole consisting of two chloridized silver electrodes separated by 0.5 cm and positioned 1 cm from the skin. The dipole delivered a 5-Hz sinusoidal amplitude modulation at approximately 20% contrast. The body surface was initially surveyed at relatively large spatial intervals to identify the responsive region; positions near its perimeter were then sampled more closely. Responses were monitored from the spike train and the corresponding peri-stimulus time histogram, with the receptive-field edge defined by positions where the ON response to the stimulus transitioned to no response (or OFF responses if driving the surround RF). The receptive field’s rostrocaudal and dorsoventral extent was recorded relative to the dimensions of the fish.

### Cerebellar feedback manipulation

Responses were first recorded with the indirect cerebellar feedback pathway intact and then after pharmacological blockade. For most neurons, a blunt glass micropipette connected to a picospritzer (Parker) was positioned in the praeminential–cerebellar tract carrying projections from the nucleus praeminentialis (nP) to the EGp. A saline solution with 2% lidocaine hydrochloride (Hospira) was pressure-ejected into the tract, with the injected volume adjusted to span the diameter of the tract while limiting spread to surrounding tissue. This manipulation prevents electrosensory activation of EGp granule cells and thereby silenced their parallel-fiber input to the ELL (Bastian 1986; Marsat and Maler 2012).

For a subset of 15 neurons, feedback was blocked locally by pressure-ejecting 1 mM disodium CNQX (Sigma-Aldrich) into the ELL molecular layer near the apical dendrites of the recorded pyramidal cell. The double-barrel injection pipette was positioned by moving it and periodically ejecting brief pulses of glutamate (1 mM); a short-latency increase in firing rate indicated proximity to the target dendrites. CNQX was then delivered as a single injection/100-ms pulses at 0.5 Hz for 20 s. This focal manipulation blocks non-NMDA glutamatergic transmission from parallel fibers, including both their excitation of pyramidal-cell apical dendrites and the disynaptic inhibition recruited through molecular-layer interneurons (Bastian 1993). Results for this method were similar to the results obtained when blocking the nP-EGp tract; therefore, we pooled the results.

The effectiveness of each manipulation was assessed from responses to a spatially broad 5-Hz beat. A block was considered successful when the normally attenuated response became more strongly phase-locked to the beat, with an increase in response gain or peak-to-trough firing-rate modulation (Bastian 1986; Bol et al. 2011; Chacron et al. 2005). Preservation of the response to a 20 Hz localized stimulus centered on the receptive field confirmed that the drug had not directly affected pyramidal cell excitability or feedforward inputs.

### Stimulation

Stimuli were generated in MATLAB (MathWorks) and sampled at 20 kHz. The fish’s electric organ discharge (EOD) was monitored with electrodes near the head and tail, and each EOD cycle triggered a DG1022A sine-wave generator (Rigol) to produce a phase-locked carrier. This signal was multiplied by the desired amplitude-modulation waveform and delivered through a Model 2200 stimulus isolator (A-M Systems), producing controlled modulations of the fish’s electric field (Benda et al. 2005). Step stimuli, broadband noise, and sinusoidal amplitude modulations were initially presented through two 30-cm carbon electrodes positioned parallel to the fish to characterize each neuron.

Experimental stimuli were delivered through an artificial conspecific, or “fish-pole,” that reproduced the spatially heterogeneous electric field of another fish (Milam and Marsat 2026). The source consisted of an asymmetric dipole formed by chloridized silver electrodes embedded in agarose and separated by approximately 4–4.5 cm. Its geometry and output were calibrated against measurements from live fish, preserving the spatial gradient and distance-dependent attenuation of natural conspecific signals rather than producing uniform stimulation across the skin.

Three stimulus types were presented: constant-contrast sinusoidal amplitude modulations (SAMs); sinusoidal-envelope modulations (SEMs), in which beat contrast varied at 0.5 Hz; and random-envelope modulations (REMs), with envelope frequencies between 0 and 0.5 Hz (i.e. flat spectrum up to 0.5 Hz, rapidly decreasing thereafter). For each of these, the carrier AM beat was either 5, 75, or 150 Hz. For one dataset (n=46 neurons), 5 Hz AM beats were presented with the fish-pole oriented perpendicular to the receiver. The fish-pole was placed 10 cm from the body in one of three locations: facing the operculum (i.e., close to the head), midbody, or facing the caudal 25% near the tail. Another dataset (n=32 neurons) used beat frequencies of 5, 75, or 150 Hz with the fish-pole positioned 10 or 25 cm from the receiver facing the mid-body.

### Data Analysis

#### Initial response quantification

All analyses were performed in MATLAB using custom scripts. Extracellular recordings were filtered and spike-sorted offline to isolate the largest consistently shaped waveform attributable to a single neuron. Spike times were converted to binary sequences sampled at 2 kHz.

Responses to beat stimuli were folded across stimulus cycles to construct peri-stimulus time histograms (PSTHs), representing spike probability as a function of beat phase. Four complementary measures were used to characterize each response. The synchronization coefficient quantified the precision of phase locking between spikes and the beat cycle, ranging from zero for no consistent phase preference to one when all spikes occurred at the same phase (Goldberg and Brown 1969). Peak-to-trough firing rate was calculated by smoothing the spike train with a sliding window equal to one-sixth of the beat period, measuring the difference between the maximum and minimum firing rates within each cycle, and averaging this difference across cycles. Beat-response gain was defined as the amplitude of a sinusoid at the beat frequency fitted to the cycle-averaged PSTH. Mean firing rate was calculated across the full duration of a response. Together, these measures captured overall activity, phase locking, and the magnitude of beat-evoked firing-rate modulation.

#### Relative response quantification (i.e., feedback cancellation)

The effect of feedback on each response measure was quantified by comparing response measures obtained with feedback intact, *R*_*FB*_, and after feedback blockade, *R*_*BLK*_. A relative response, *ReR*, in percent was calculated as:

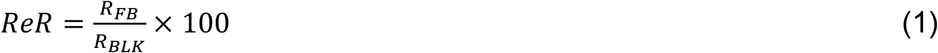

A relative response of 100% indicated the same value of the response measure with feedback present and blocked. Values below 100% indicated attenuation by feedback, whereas values above 100% indicated a larger value with feedback present. For example, a relative response of 80% corresponded to a 20% reduction relative to the feedback-blocked condition. Note that in some cases, a weak response when the feedback is blocked can become a stronger response when the feedback “over-cancels” it: the peak is attenuated to 0 and the response at the trough is increased to cause a larger peak in firing rate. Relative response was calculated separately for synchronization, peak-to-trough firing-rate modulation, response gain, and mean firing rate.

#### White noise envelope (REM) stimuli analysis

Instantaneous firing rates of responses to REM stimuli were calculated from binarized spike trains by convolving them with a Gaussian filter (width equal to 1/6^th^ the AM carrier period). Linear encoding of the envelope was quantified for each feedback condition using the magnitude-squared coherence between the envelope, *E*(*t*), and instantaneous firing rate, *R*(*t*):

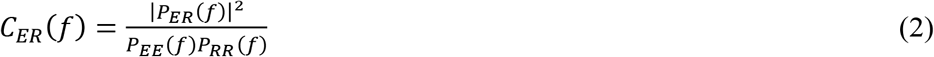

where *P*_*ER*_ is their cross-spectral density and *P*_*EE*_ and *P*_*RR*_ are their respective power spectral densities. Coherence ranges from zero to one and indicates the fidelity with which firing-rate fluctuations linearly followed the envelope at each frequency within its 0–0.5-Hz bandwidth (Borst and Theunissen 1999).

To determine how recent envelope history shaped the feedback effect, we first smoothed the firing rate estimates with a 200 ms boxcar filter to obtain a measure that reflects mean firing rate over at least one AM cycle while preserving envelope-frequency modulations. We calculated the firing-rate difference as the smoothed firing rate after feedback blockade minus that with feedback intact. This difference, expressed in spikes/s, was used for the kernel analysis (note that it is distinct from relative response in Equation 1).

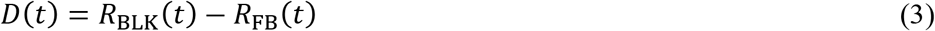

Positive values of *D*(*t*) therefore indicated feedback-dependent attenuation of the envelope response. After subtracting the means of *D*(*t*) and *E*(*t*), a linear kernel relating preceding envelope amplitude to the subsequent feedback effect was calculated in the frequency domain as

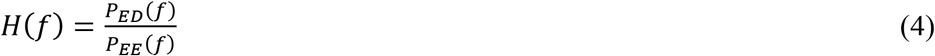

and transformed into the time domain using the inverse Fourier transform. The resulting kernel describes how envelope amplitude at each preceding time lag contributed to the current difference between the feedback-intact and blocked responses. Positive kernel values indicate that stronger preceding stimulation predicted greater subsequent attenuation by feedback. Statistical confidence limits were obtained by flipping the time sequence of the response relative to the envelope, recalculating the kernel for each shuffled dataset, and using two times the standard deviation as the limit.

#### Population decoding of spatial information

Spatial coding was quantified using a weighted Euclidean distance analysis (for more details, see: Marsat et al. 2023; Milam and Marsat 2026). Responses were divided into 1-s epochs, and synchronization coefficient, peak-to-trough firing-rate modulation, beat gain, and mean firing rate were calculated for each epoch. Each measure was analyzed separately. For a given pair of stimulus locations, pseudo-population responses were assembled by selecting one epoch from each of *n* neurons and representing the resulting response as a point in an *nn*-dimensional space, with each dimension corresponding to one neuron. Analyses were performed independently with feedback intact and blocked.

For each neuron *n* and stimulus pair (e.g., location *M* and *T*), the responses’ dimensions was weighted based on the area difference between the two location-specific response distributions *P*:

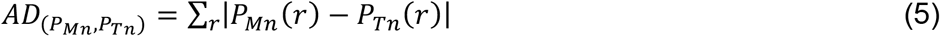

from which the weights *W* were calculated by normalizing to 1 across the population of neurons:

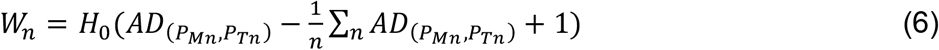

where H_0_ is the Heaviside step function.

Weighted Euclidean distances were then calculated between population responses elicited by the same location, *P*(*D*_*xx*_), and by the two different locations, *P*(*D*_*xy*_). The separation of these distributions was quantified using a receiver operating characteristic analysis. For each distance threshold *T*,

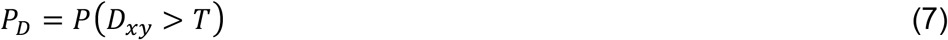

and

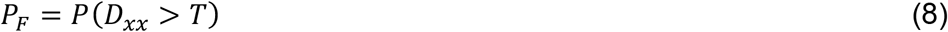

where *P*_*D*_ and *P*_*F*_ are the probabilities of correct and false discrimination, respectively. The discrimination error was calculated as

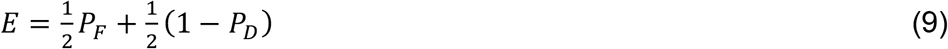

and the minimum value obtained across thresholds was retained. Error values therefore ranged from 0.5 for chance-level discrimination to 0 for complete separation of the population responses.

The analysis was repeated while varying the number of neurons included in the pseudopopulation. For the three rostrocaudal stimulus positions, decoding was performed for the head-middle, middle-tail, and head-tail comparisons and then averaged across pairs. The decrease in error with population size, *n*, was fitted with:

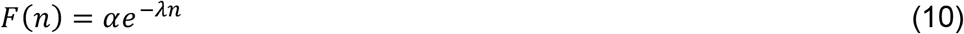

where *λ* provides a measure of population coding efficiency: larger values indicate that each additional neuron contributes more information about stimulus location.

#### Statistical analysis

Statistical analyses were performed in MATLAB or in R. Data distributions were tested for normality with a Shapiro-Wilk or a Kolmogorov–Smirnov test before further analysis, and the corresponding parametric/non-parametric analyses were used as specified in the results. All tests were two-tailed unless otherwise stated, and statistical significance was defined as p<0.05. The statistical test, sample size, and p-value associated with each analysis are provided in the corresponding figure legend when p is below the significance threshold.

## Results

Our experimental approach replicates realistic social encounters by recording in vivo in awake fish and creating conspecific signals with realistic spatial structures and natural temporal modulations. The stimulus dipole used to generate conspecific signals, labeled the “fishpole”, has been calibrated in shape and intensity to mimic an average-sized conspecific (see Methods). To keep the stimulation protocol short and keep it within the effective duration of the pharmacological feedback block, we collected two datasets in which we varied two aspects of the stimulus. In the first set, we focused on varying the beat frequency; in the second dataset, we varied the spatial aspect (fishpole position) (Figure 1A).

**Figure 1.**
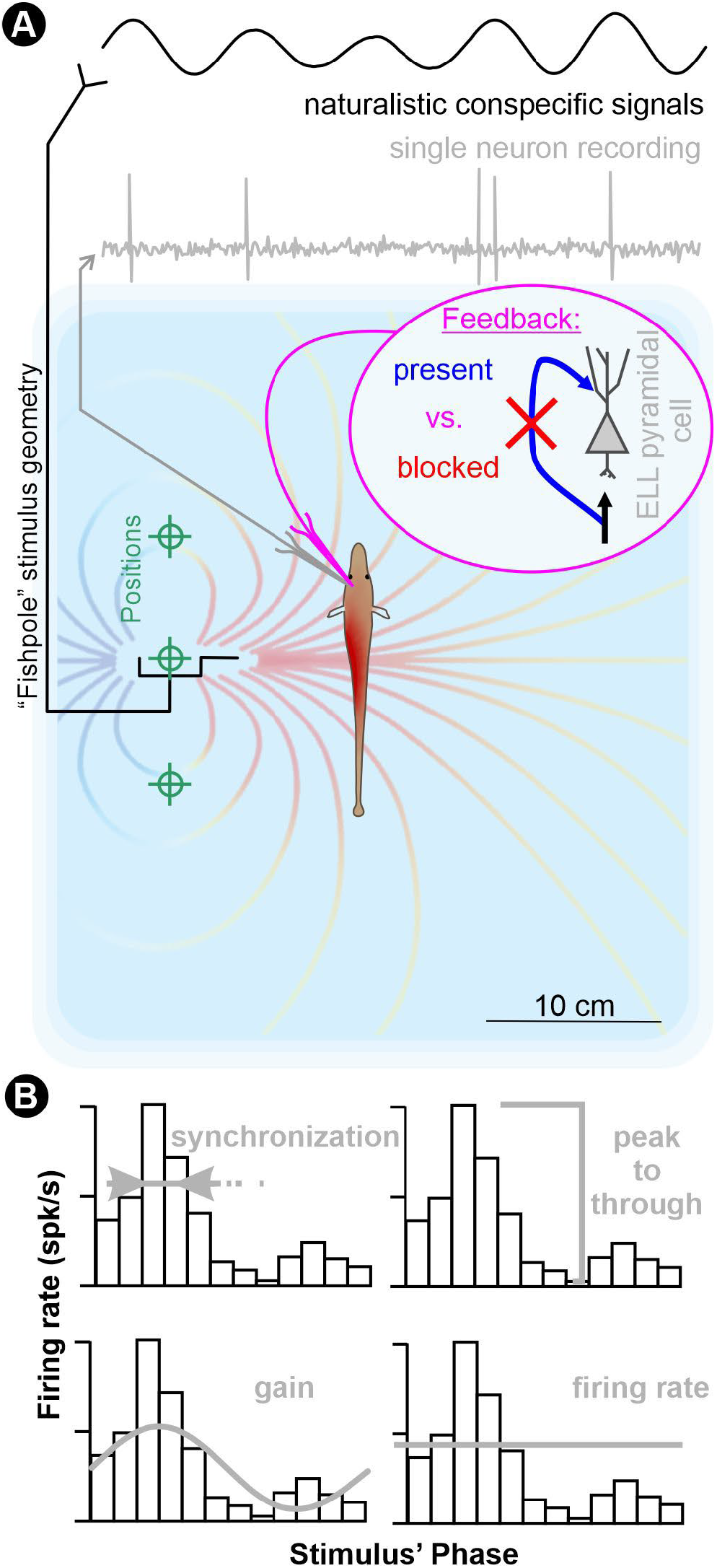
Illustration of the experimental design. **A.** The experimental setup consists of a stimulation dipole replicating the spatial structure of conspecific EODs (fishpole, black lines) and that stimulates the focal fish with a realistic modulated beat signal (top black trace). Field lines of this signal are represented qualitatively with the blue-to-red background pattern. The stimulation pattern onto the focal fish’s body is represented here with a red hotspot on the side facing the depicted fishpole. The fishpole is positioned 10 cm away (unless otherwise noted) at three different positions (green marks) near the head, mid-body, or tail. Single-cell recording in the awake focal fish is performed (grey lines) while another micro-pipette allows to pharmacologically block the feedback (magenta inset). **B**. Neural recordings are initially characterized by their response to the beat of the stimulus. A PSTH of the firing relative to the beat phase allows to quantify the synchronization coefficient, the peak-to-though firing rate, the response gain at the beat frequency, and the mean firing rate of the response. These quantifications are summarized graphically with the grey illustrations onto the black PSTHs.

### Feedback cancellation of constant beat AM signals

We first examine the effect of feedback on the responses to the simplest signals: a constant beat of a given AM frequency with no envelope modulation. This allows for a direct comparison with previous research. In particular, we aim to confirm that the qualitative effect of feedback cancellation when spatially realistic conspecific signals are used (delivered through our fishpole) is comparable to the effects described previously for spatially uniform stimuli using a global dipole configuration (large electrodes positioned on each side of the fish; e.g. Bol et al. 2011).

The response strength for each neural recording was quantified through 4 measures that assess both the number of spikes and the spiking pattern relative to the stimulus (Figure 1B). The synchronization coefficient reflects the spike patterning: how much the spikes are clustered around a specific phase of the beat AM (e.g., a neuron firing only, and very precisely, at the top of the beat would have a response synchronization coefficient of 1). The mean firing rate, on the other hand, is unaffected by the spiking pattern and timing relative to the beat’s phase. Response gain and peak-to-trough firing rate reflect both the firing strength and timing: Neurons that fire more and in phase with the stimulus will have higher response gain and peak-to-trough firing rate.

We quantify the effect of feedback as relative response: 100% indicates equal response measures with feedback present and blocked. For example, synchronization values of 0.4 with feedback present and 0.5 with feedback blocked therefore correspond to an 80% relative response (i.e., a 20% reduction due to feedback). We find a marked cancellation of the low-frequency beat (5 Hz) where the feedback causes a decrease in the response at the peak of the beat and an increase at the trough (Figure 2). This change in spiking pattern is reflected in relative responses below 100% for gain, peak-to-trough firing rate, and synchronization. Mean firing rate was not significantly affected by the feedback. These results are in line with previous findings and can be explained by the fact that feedback inputs include both an inhibitory and an excitatory component. The feedback input is thereby a mirror image (in terms of phase relationship) of the feedforward input without a strong excitatory or inhibitory overall effect.

**Figure 2.**
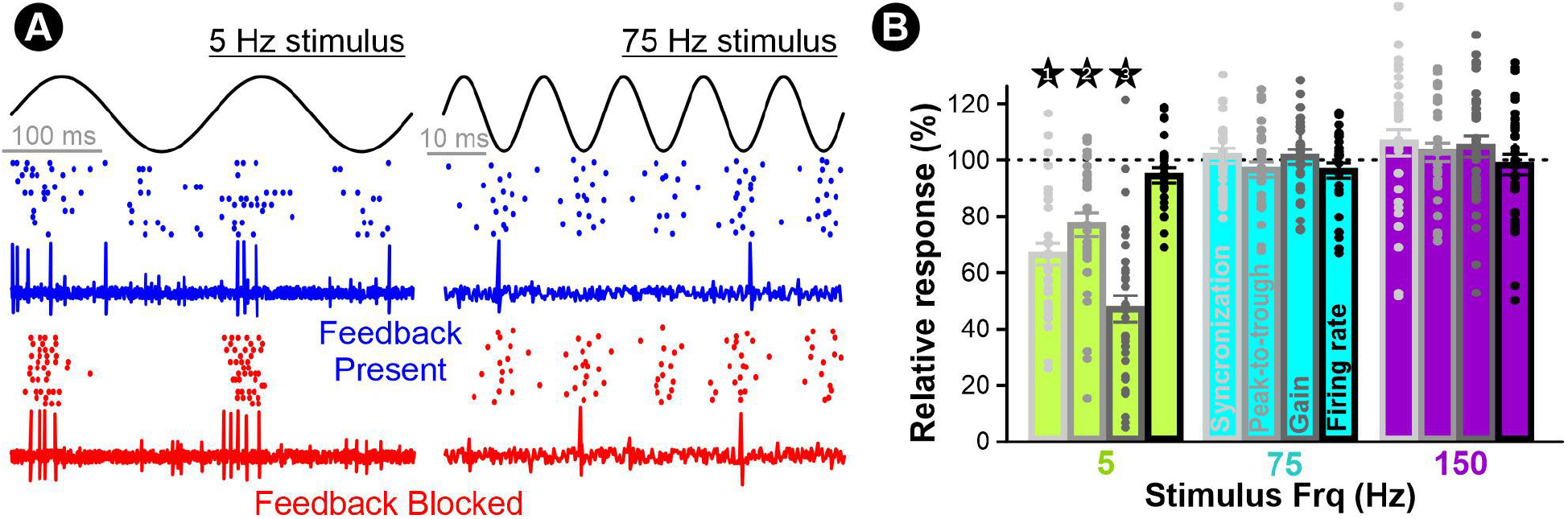
Basic impact of feedback on responses to constant beats. **A.** Example of a neuron’s response to two beat frequencies (low: 5 Hz and high: 75 Hz) with the feedback pathway blocked (red) or not (blue). A few cycles of the beat stimulus are shown along with sample traces of extracellular recordings and raster plots of the neuron’s firing pattern. **B**. Mean (± s.e.; n=32 neurons) response strength when the feedback is present relative to the response when feedback is blocked for constant beats signals. The dotted line at 100% indicates no difference between conditions; values below 100% indicate attenuation by feedback, and values above 100% indicate a larger response measure with feedback present. Relative responses were tested against the no-difference value of 100% using a one-sample t-test with Holm correction for multiple comparisons. P-values below 0.05 are indicated by numbered asterisks: *1, p<10^−8^; *2, p<10^−4^; *3, p<10^−11^. Note that the firing rate for 5 Hz stimuli might have been slightly affected by the feedback: the p-value is 0.02 before Holm correction and 0.08 with correction.

Also confirming previous findings, we show that the pattern of responses to constant beat signals of high frequency is not affected in a beat-phase-specific manner, as none of the response measures are significantly different when the feedback is blocked. Overall, our relative response of ∼50% for gain at low frequencies (a ∼50% reduction) and ∼100% at higher frequencies (little or no reduction) is qualitatively similar to previous research finding strong reduction of the response (decrease of 60%-80%) for frequencies below 15-20HZ but no cancellation for higher frequencies (Bol et al. 2011). We do not provide an exact quantitative comparison because it would require a stimulation protocol that adjusts the intensity of the uniform-global vs fishpole stimuli within the receptive field of the cell being recorded. This question, and the corresponding set of experiments, is beyond the scope of the present study. Nevertheless, our results confirm that feedback has a marked canceling impact on low-frequency responses to spatially realistic conspecific signals, albeit possibly weaker than for stimuli configurations with a more global and uniform spatial distribution.

### Cancellation of dynamic AMs

The beat signals that arise from social interactions with another fish will commonly undergo additional modulations due to the relative movement of the two fish. As fish get closer, the contrast of the beat will increase from being imperceptible (0% contrast -i.e., no beat-if the fish are very far away) to modulating the focal fish’s EOD 100% or more when they are in very close proximity (i.e., the combined signal would be twice the focal fish’s baseline EOD when the two fish’s EODs are in phase, and close to null when they are in antiphase). These movement-related contrast envelopes consist of very low-frequency modulations related to the speed of movement that typically contain most power <1 Hz (Fotowat et al. 2013). The cerebellar feedback dynamically adapts to the signal’s properties -including signal strength-through synaptic plasticity (Bol et al. 2011; Harvey-Girard et al. 2010; Lewis and Maler 2004) and possibly other circuit-level mechanisms (Johnson et al. 2025). Therefore, we expect the effect of feedback to be influenced by the change in contrast caused by the fish’s relative movements. To examine the impact of contrast envelopes on beat cancellation, we use sinusoidal envelope signals because they allow the most controlled and repeated signal pattern and analysis. The corresponding behavior would amount to a pair of fish quickly moving back and forth towards each other in a smooth and periodic manner. Although this movement pattern might be too regular to be naturally observed frequently and produced for many cycles, the stimulus encapsulates the relevant temporal properties. The signals we use thus consist of beats of low, medium, or high frequencies (5, 75, 150 Hz) modulated in contrast (0 to 100%) at a frequency of 0.5 Hz (Figure 3).

**Figure 3.**
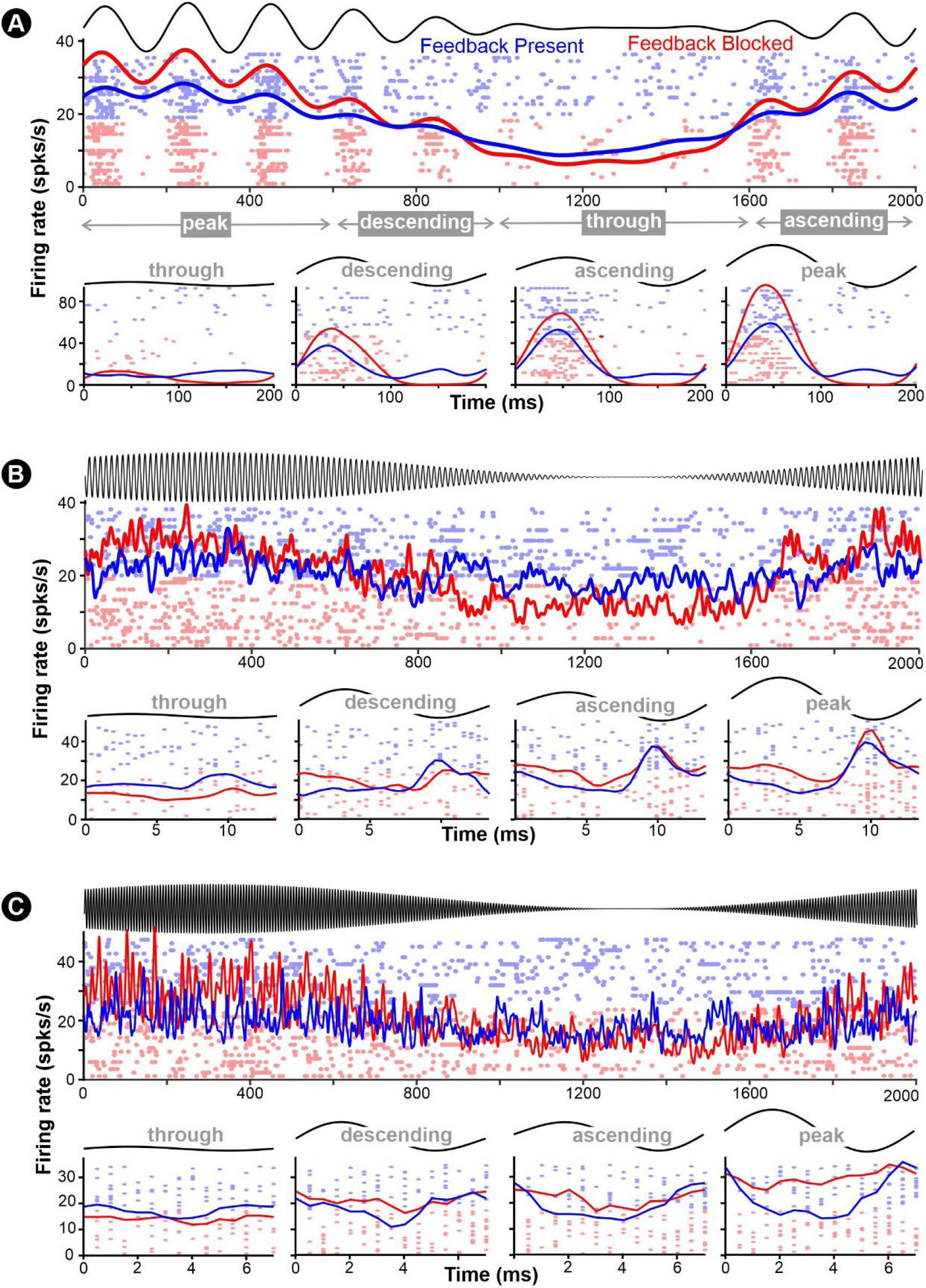
Responses to contrast-modulated beat depend on envelope phase. In each panel we show the stimulus in black and an example of a population response in blue (feedback present) and red (feedback blocked). The responses display a raster plot with a randomly chosen subset of neurons and excerpts. Overlaid on the raster plots is a population firing rate for the excerpt. The beat frequency is either low (5 Hz, panel **A**) or high (75 Hz, panel **B**; 150 Hz, panel **C**). The upper half of each panel shows the response to one cycle of the 0.5 Hz contrast envelope. The bottom half shows the responses to one beat cycle, where each of the four plots represents a beat cycle at different phases of the envelope (trough, descending phase, ascending phase, or the peak of the envelope cycle).

As expected, the firing rate and beat-synchronization of the responses increase with contrast. For high-frequency beats, most cells do not phase-lock well to the beat cycles; nevertheless, firing rate increases with contrast. Comparing the responses with and without feedback for low-frequency beats, we clearly see the beat-phase-specific effect of feedback. For example, looking at spiking patterns in the descending phase of the envelope (Figure 3A), we see an increase in firing at the trough of the beat and a decrease in firing at the peak due to feedback. This effect, however, is less pronounced for similar-intensity beat cycles in the ascending phase of the envelope. This suggests that the strength of the feedback is influenced by the strength of the signal in recent history. In the ascending phase of the envelope, recent beat cycles were relatively weaker compared to the current cycle, and the opposite is true for the descending phase of the envelope.

At higher beat frequency, even though the feedback does not provide a mirror image cancellation of the peak-and-trough response to the beat cycle, it does influence the mean firing rate in an envelope-phase-specific manner (Figure 3B-C).

To confirm this qualitative observation, we quantify in Figure 4 the beat-canceling effect of feedback on 4 aspects of the response (from synchronization to mean firing rate) as a function of the envelope phase. Although the response timing relative to the beat is only significantly affected for low-frequency beats (Figure 4A), the overall firing rate is significantly different for the envelope’s peaks and troughs between responses for which the feedback is present or blocked (Figure 4D). Indeed, the feedback attenuates the firing rate at the peak of the envelope (relative response<100%) and increases firing rate at the trough of the envelope cycle (relative response>100%). These results point out that not only is the effect of feedback on beat-coding influenced by the recent strength of the signal, but feedback also affects how mean firing rate encodes envelope modulations.

**Figure 4.**
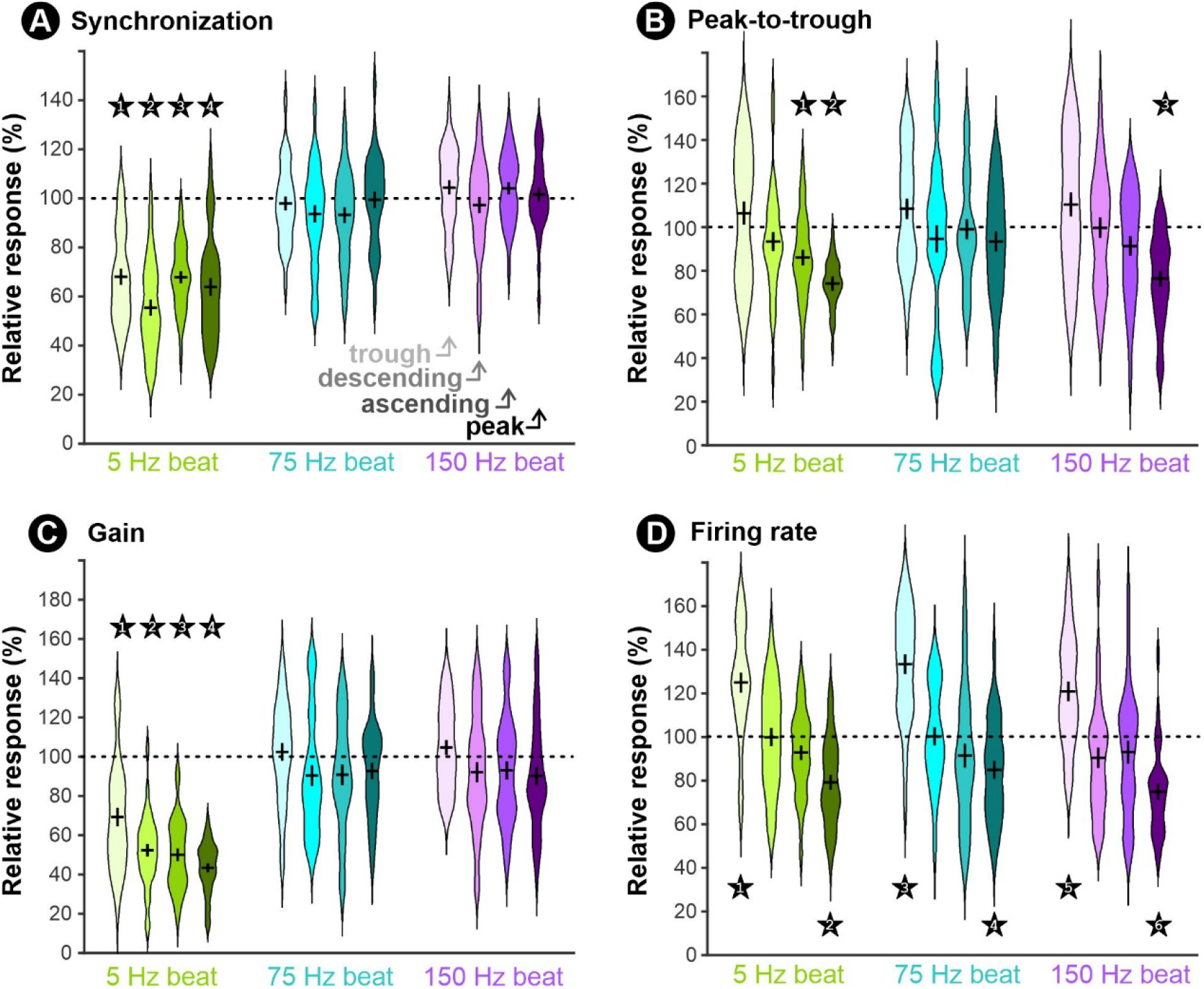
Cancellation of beat response is dependent on envelope modulations. Relative response is shown for different envelope phases (shades; see Figure 3) and beat frequencies (colors). The dotted 100% line indicates no difference between feedback conditions; lower and higher values indicate smaller and larger responses with feedback present. The response strength was measured based on one of the four basic response characterizations: **A**. Synchronization. **B**. Peak-to-trough firing rate. **C**. Response gain. **D**. Mean firing rate. Violin plots show the distribution of relative responses across neurons (n=32), with the mean marked by the horizontal bar of the cross and the standard error by the extent of the vertical bar. Statistical differences between responses with feedback and without feedback were tested with a Scheirer–Ray–Hare test in each panel (different measures) and for each frequency independently (with envelope phase and feedback/no-feedback as the two factors). Results below the 0.05 significance threshold for the feedback factor or feedback-phase interaction were then analyzed pairwise using Dunn’s tests with Holm correction for multiple comparisons. Significant values are indicated by numbered asterisks with exact values as follows. **A**: *1, p<10^−9^; *2, p<10^−12^;*3, p<10^−12^; *4, p<10^−9^. **B**: *1, p=0.0036; *2, p<10^−12^;*3, p=5·10^−6^. **C**: *1, p=4·10^−6^; *2, p<10^−13^;*3, p<10^−14^; *4, p<10^−20^. **D**: *1, p=5·10^−5^; *2, p=2·10^−5^;*3, p<10^−6^; *4, p=0.007;*5, p=4·10^−4^; *6, p<10^−6^.

### Contrast envelope cancellation

The contrast envelope used for the results in Figure 3 consists of sinusoidal modulation, allowing us to compare the canceling effect of feedback using the same simple response measure as for the beat. Figure 5 shows relative responses based on overall gain at the beat frequency and mean firing rate, summarizing the feedback’s canceling effects described in the previous section. We also calculate relative response from the gain at the envelope frequency (0.5 Hz), which measures how strongly firing-rate modulations follow the contrast envelope. Our results confirm that the response to envelopes is canceled by feedback as the gain is reduced by ∼50%. Notably, this envelope cancellation is comparable irrespective of the beat frequency (repeated-measures one-way ANOVA, n=32, p=0.64). Together, these results imply that feedback shapes the coding of social movement-related envelopes across a broad range of naturalistic conditions extending beyond interactions between fish with similar EOD frequencies.

**Figure 5.**
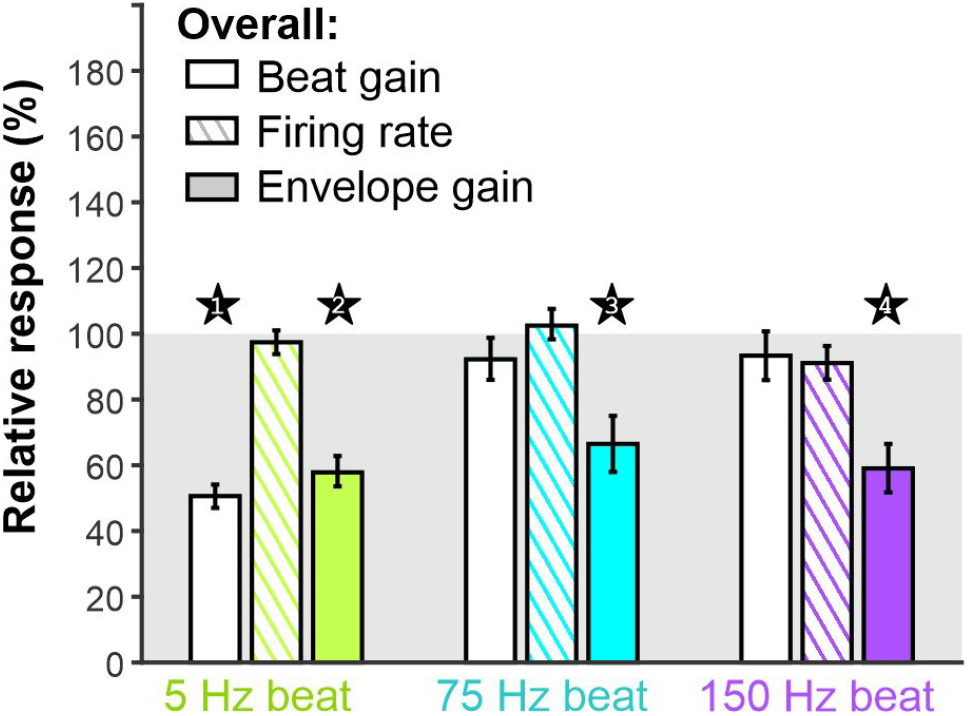
Feedback affects the coding of envelope modulation for both low and high beat frequencies. Using the same envelope-modulated beat signals as in Figure 3-4, we quantified three aspects of the overall response to these signals: the overall response gain at the beat frequency, the overall response gain at the envelope frequency, and the mean firing rate. For each measure, bars show the mean relative response (± s.e.; n=32 neurons), calculated using Equation 1. Relative responses were tested against the no-difference value of 100% using a one-sample t-test with Holm correction for multiple comparisons. P-values below 0.05 are indicated by numbered asterisks: *1, p<10^−15^; *2, p<10^−8^; *3, p=0.0005; *4, p<10^−5^.

To further examine this effect, we created stimuli with a more irregular envelope modulation consisting of random modulation (0Hz to 0.5 Hz Gaussian white noise). These stimuli have the advantage of allowing specific analytical approaches because they do not have the long-range temporal correlations of sinusoidal envelopes. They also reproduce the more stochastic envelope modulations caused by variable movement patterns. These stimuli were created to have an envelope standard deviation of roughly 30% contrast (and thus only the largest peaks in contrast are close to 100%) and thus mimic more moderate movement amplitudes compared to our sinusoidal envelopes that reached 100% contrast for each envelope peak. We show in Figure 6A an excerpt of these stimuli (here with a low-frequency beat) showing the advantage of the random envelopes and the reduced temporal correlations in beat amplitude. Specifically, we point out three beat cycles showing the range of relationships between a given cycle’s amplitude and the amplitude of previous cycles. In panel B, we focus on a strong beat cycle preceded by several very weak beat cycles. In contrast, in panel C, a similarly strong beat cycle is preceded by similarly strong cycles. Given the random nature of the envelope, we have a wide range of cycle/previous cycles relationship such as the additional example in panel D, where a weak cycle is preceded by strong beat cycles. Qualitative observation of the response patterns to these three examples shows the same effects that were observed for sinusoidal envelopes, where the strength of the feedback effect is linked to the amplitude of the envelope on preceding cycles. Compare, for example, panels B and C (Figure 6), where the cancellation is more pronounced when the previous beat cycles are stronger (C). Furthermore, firing rate modulations and how they follow envelope modulations are attenuated by feedback, as shown by the reduced coherence between envelope and firing rate (Figure 7A). This envelope cancellation is comparable for all three beat frequencies tested.

**Figure 6.**
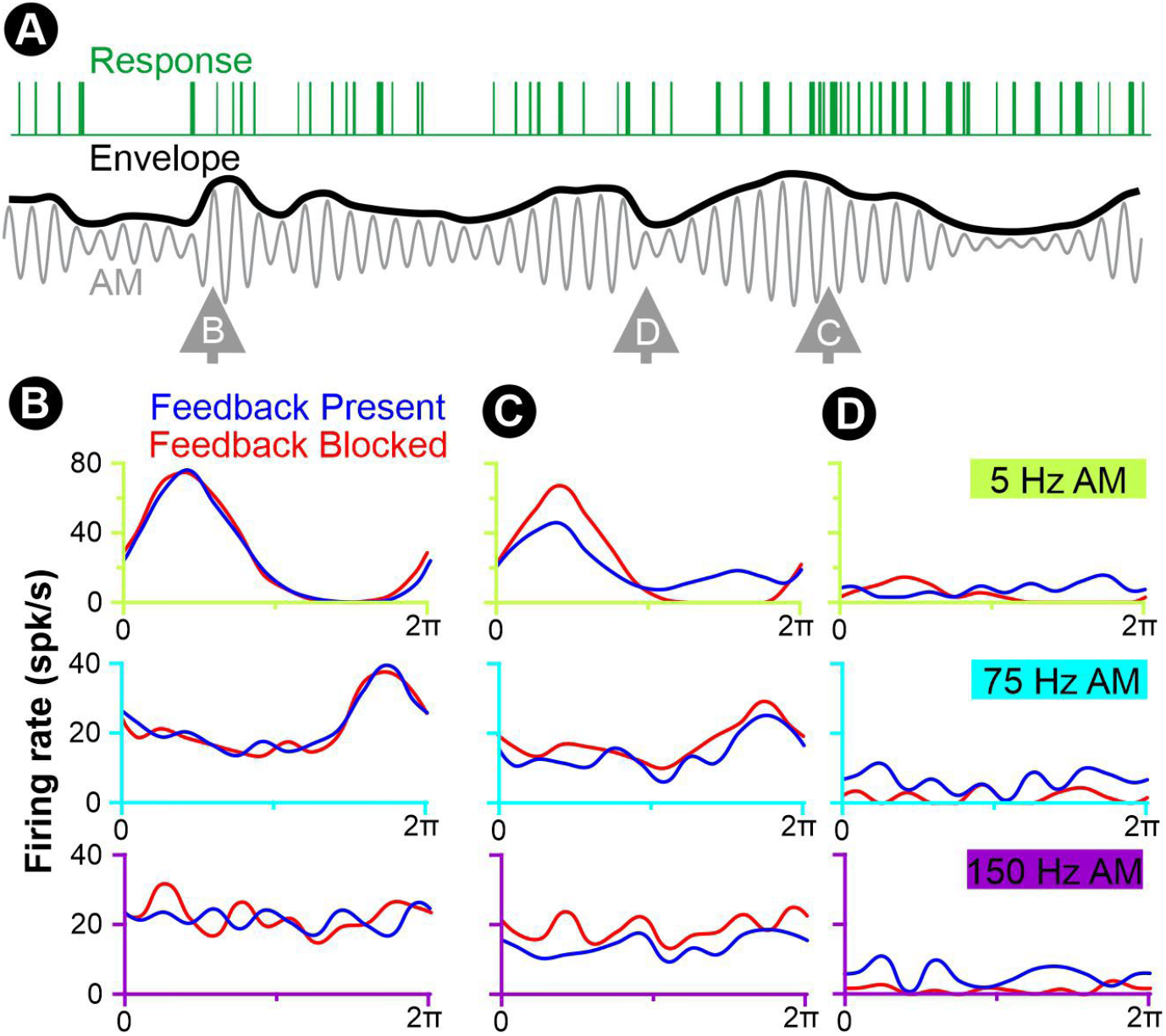
Responses to random envelope modulations show that beat cancellation depends on recent contrast strength (envelope). **A.** Example of a binarized response (green) to a 5 Hz beat (grey) modulated with a random contrast envelope (0 to 0.5 Hz; black). We selected three specific cycles to illustrate the effect of recent beat contrast on cancellation: **B**. Strong beat cycle preceded by weak ones. **C**. Weak beat cycle preceded by strong ones. **D**. Strong beat cycle preceded by strong ones. For each of these example time points relative to the envelope, we show the mean response (n=32) to the beat cycle for low (5Hz, top plots) and high (75 or 150 Hz, bottom plots) beat frequencies. We compare responses when the feedback is present or blocked (blue and red firing rate traces, respectively)

**Figure 7.**
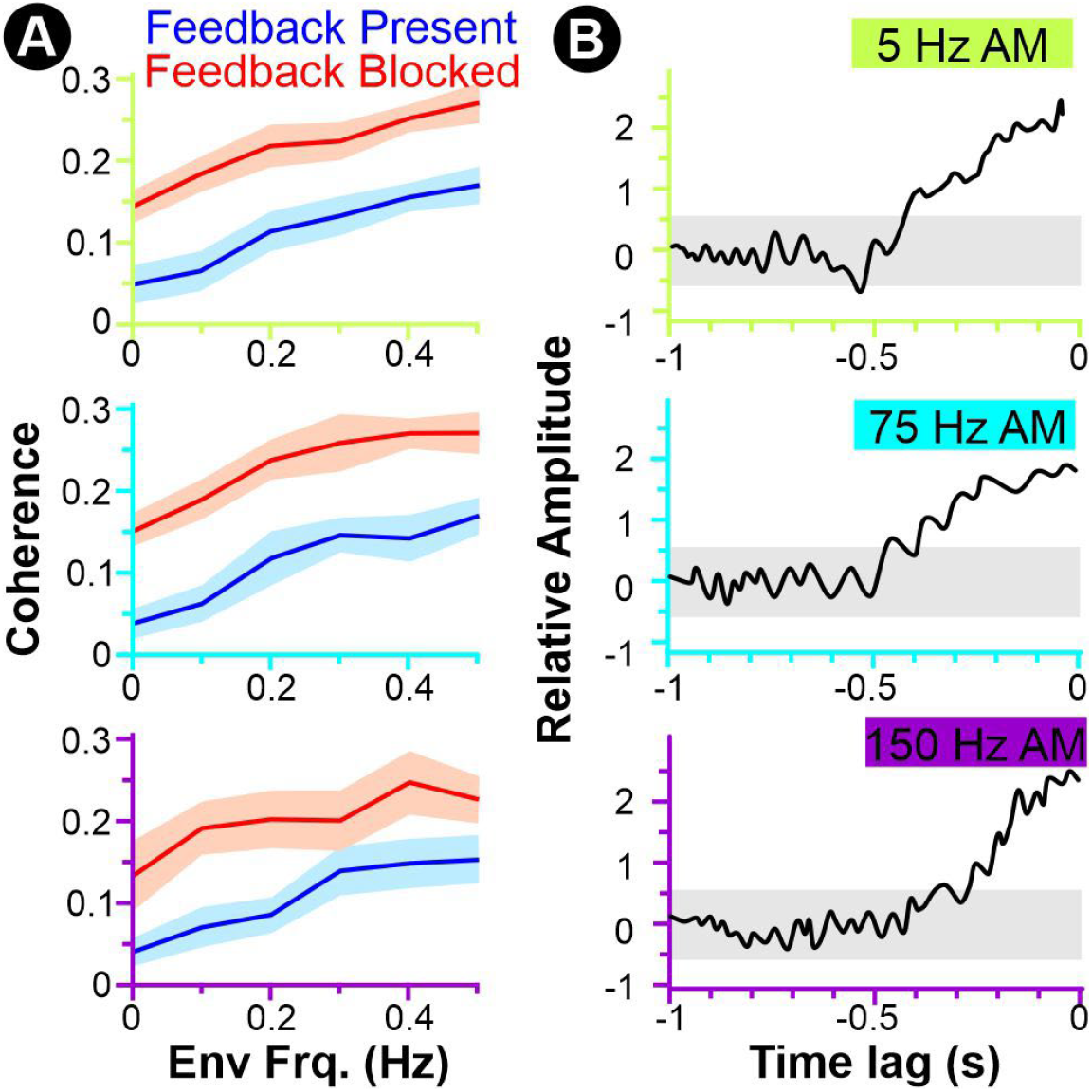
The correlation between the envelope and responses’ firing rate is attenuated by feedback according to a linear function of the recent beat contrast. **A.** Mean coherence (± s.d.; n=32 neurons) between the firing rate and the envelope. While the random envelope modulation is the same in each panel, the beat frequency is: 5, 75, or 150 Hz (from top to bottom). The differences between the coherence when feedback is present (blue) or blocked (red) are significant in each panel (paired cluster-based permutation test, 100,000 permutations, p_FWER_<10^−5^, for all AM frequencies). **B**. We computed the linear filter relating the stimulus envelope to the population-averaged firing-rate difference (feedback blocked minus feedback present. The result can be described as a cross-correlation normalized to account for the stimulus’ autocorrelation. The grey shadow in the background shows the confidence interval defined as two times the standard deviation of the filter calculated on shuffled responses.

Random envelope stimuli also allow us to relate the feedback effect to contrast in preceding beat cycles. We calculate the linear kernel relating the envelope to the firing-rate difference (feedback blocked minus feedback present). These kernels (Figure 7B) are cross-correlations normalized by the autocorrelations in the stimulus, thereby accounting for the fact that the stimulus is correlated to itself over the time scale of its low-frequency components. As expected, the feedback strength is positively correlated with the contrast of the stimulus in recent history. The correlation extends for a 200-400 ms window preceding the current beat cycle. The properties of this filtering function provide invaluable information to characterize and reproduce the network dynamics of this feedback pathway.

### Spatial coding and background suppression

While it is important to understand the temporal dynamics of the feedback and how it affects the response properties of a given neuron, we strive to understand sensory processing with a strong focus on how it supports natural behaviors. When encountering another fish, and thus when sensing the movement-modulated beat signals, the most obvious task that the fish must accomplish is to localize the signal and possibly navigate relative to its origin (e.g., move towards or away from the other fish). While the changes in contrast of the beat can provide some cues for the distance of the other fish, it cannot guide directed swimming responses. Rather, it is the spatial distribution of the signal across the focal fish’s body that will permit localization by comparing the signal strength across receptive fields of the sensory surface.

Parallel-fiber feedback onto ELL pyramidal cells is spatially diffuse, in contrast to the more spatially restricted feedforward receptor input. Indeed, the receptive field of the main feedforward inputs typically spans 1/10^th^ to ¼ of the body’s rostro-caudal extent for neurons of the lateral and centro-lateral segments of the ELL (Milam and Marsat 2026).

The feedback onto a given pyramidal cell, on the other hand, is driven by wider bilateral receptive fields with a complex projection pattern (Chacron et al. 2005; Sas and Maler 1983, 1987), some of the cells in this pathway having inputs from almost the full rostro-caudal extent (e.g., ovoid cells of the nucleus praeeminentialis). The spatially specific feedforward input is therefore combined with a feedback input that could reflect the average sensory signal across the length of the body. This feedback is therefore adequately structured to enhance the spatial representation of the stimulus across the population of pyramidal cells, possibly by implementing a sort of background suppression, or a similar type of contrast enhancement.

We examine the effect of feedback on spatial coding by first quantifying the accuracy with which population responses in the ELL support the discrimination of stimuli from different locations. We focused this analysis on the responses to low-frequency beats since feedback affects both the firing rate and firing pattern (e.g., beat synchronization), but preliminary data suggest similar effects using higher-frequency beats (data not shown). We compare population responses using a weighted Euclidean distance approach (Marsat et al. 2023). By quantifying the separation/overlap in response patterns to different stimuli (Figure 8A), we can estimate the error probability of an ideal-observer-type decoder using ROC analysis (see Methods for details). Increasing the number of neurons included in this analysis of population responses improves discrimination accuracy because each neuron contributes some information about the stimulus (here, its location), as illustrated by the curve in Figure 8B. Therefore, we use the slope of an exponential fit of this curve to summarize the coding efficiency of the neural population (Figure 8C). A higher coding efficiency reflects the fact that a smaller number of neurons provide sufficient information for the accurate discrimination of stimuli from different locations. We find that coding efficiency is significantly reduced when the feedback is blocked (Figure 9A). This difference is visible when considering any of the four response measures we use (synchronization, gain, etc.). It is particularly pronounced when using the mean firing rate, leading to a more than two-fold difference in coding efficiency.

**Figure 8.**
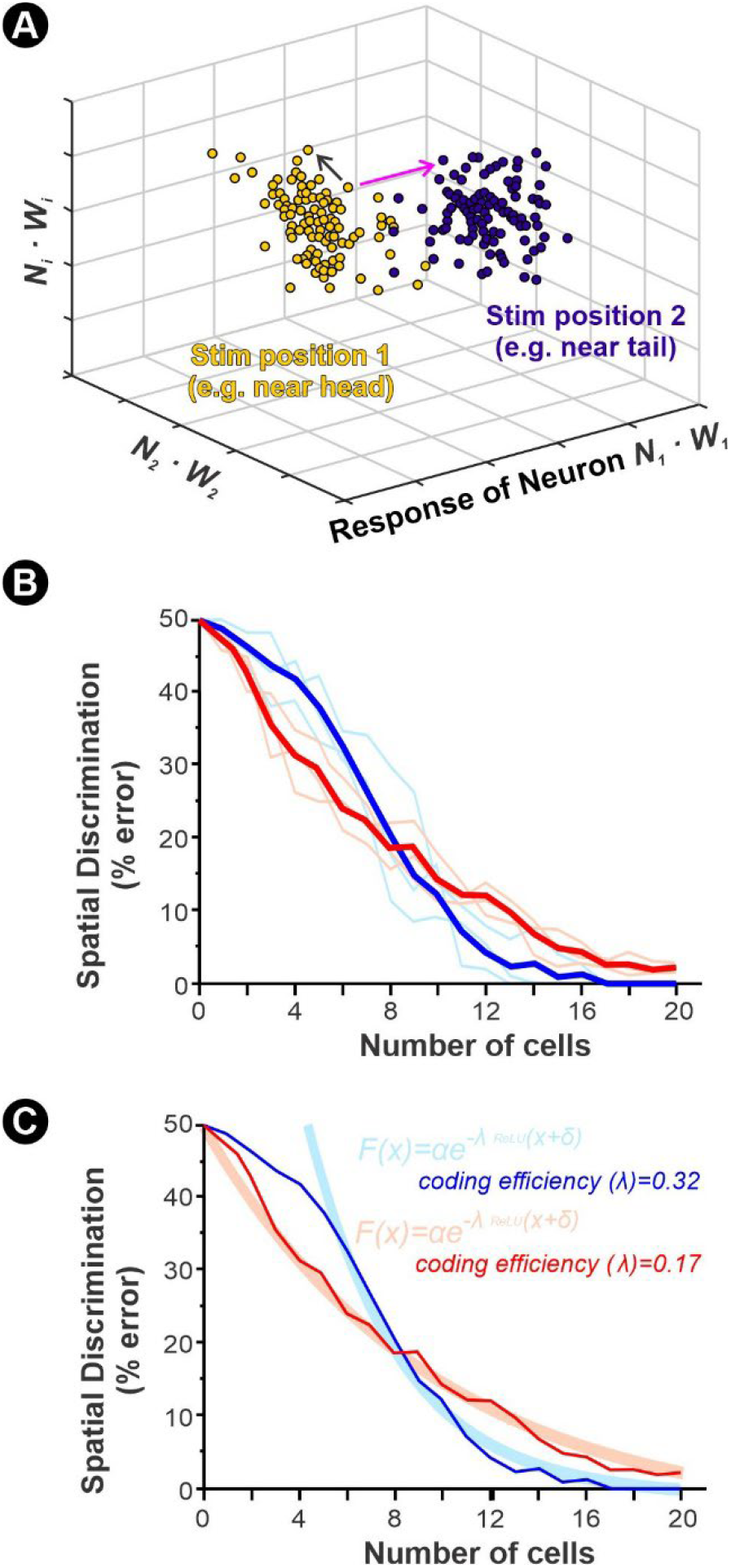
Characterizing spatial coding by comparing population response patterns for different stimulus locations. **A.** Using one of the basic measures described in Figure 1B, each neuron’s response during a 1s window is weighted by how different its response is to the two stimulus locations (i.e. how informative it is about location; see Methods for details) and constitutes one dimension of the population response. A set of population responses to position 1 (yellow) and 2 (indigo) is illustrated here in a 3D space (i.e. population considered = 3 neurons). The separation between the clusters of population responses is then characterized by taking the vector lengths between pairs of responses to the same stimulus location (black arrow) or different locations (magenta arrow). The overlap in the distributions of these vector lengths indicates how distinct the population response patterns are. An ROC analysis is used on these vector-length distributions to determine the minimum error rate that an ideal observer would make discriminating the responses for the two locations. **B**. Spatial discrimination error rate is plotted as a function of the size of the neural population used for the analysis. A larger population will allow more accurate discrimination since each neuron contributes additional information about stimulus location. The discrimination analysis is performed on pairwise stimulus locations (thin lines) and averaged across pairs (thick lines). The example shown here is based on neural responses characterized by their mean firing rates. **C**. The slope of the decrease in discrimination error as the population size is increased reflects the average information contribution of each neuron. It is therefore used to describe the coding efficiency of the population. An exponential function is fitted through these negative slopes, and the coefficient lambda (see equation inset) reflecting the steepness of this slope is therefore used to reflect the coding efficiency.

**Figure 9.**
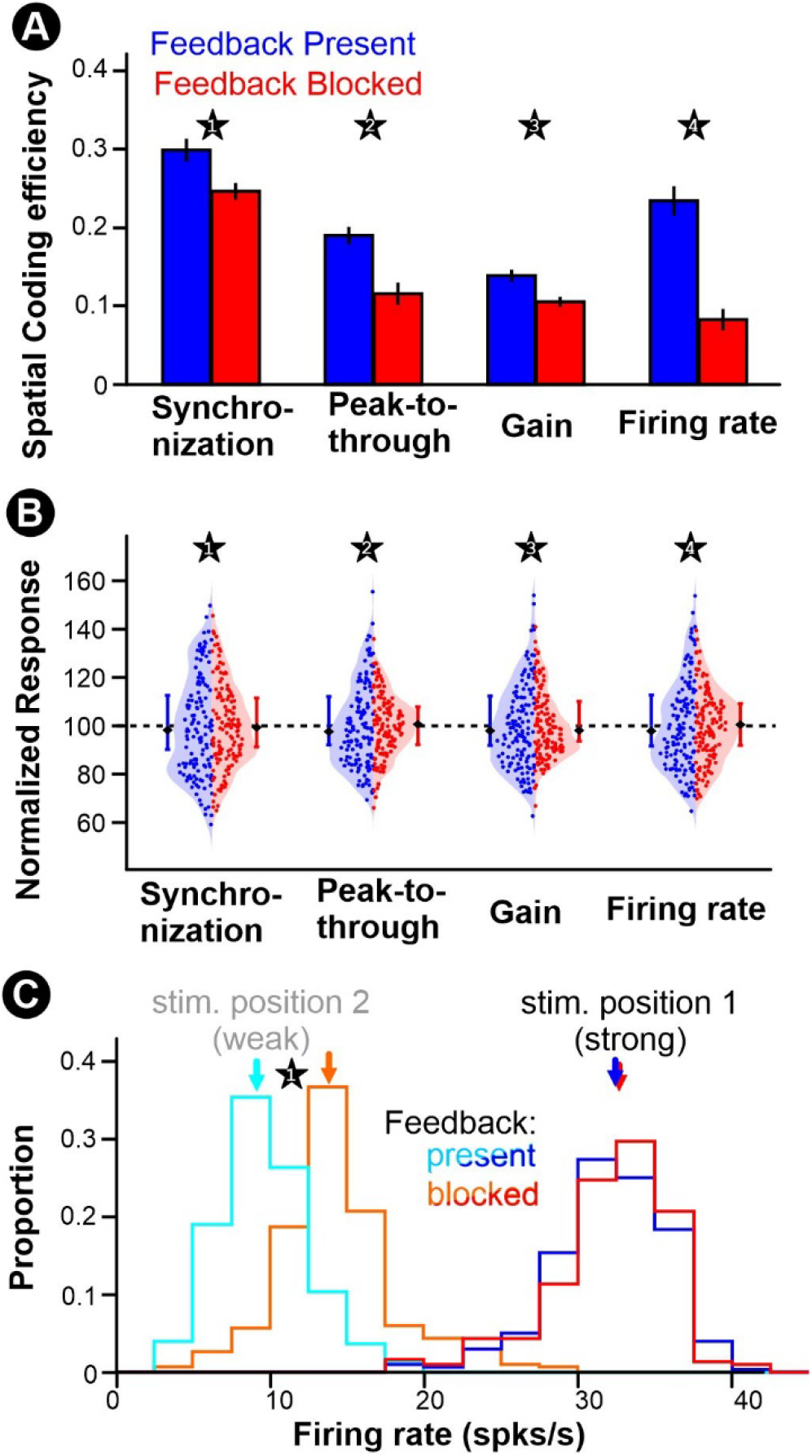
Feedback improves spatial coding by suppressing background responses and thus enhancing spatial contrast. **A.** Mean spatial coding efficiency (±s.e.) is decreased when feedback is blocked for the four response measures used. The analysis is based on recordings from 46 neurons that we used in random subsets of 20 neurons; 25 such subsets were tested. In each case, three stimulus locations were tested, leading to 3 stimulus pairs. We averaged the spatial discrimination curves across the 3 stimulus pairs to obtain a single coding efficiency for a given population subset. Therefore, the sample size used for displaying the variability of the data and for statistical analysis is n=25. We find a statistically significant decrease in coding efficiency when feedback is blocked (Wilcoxon test); significant values are indicated by numbered stars with exact values as follows: *1, p=0.005; *2, p=0.004; *3, p=3·10^−4^; *4, p=5·10^−5^. **B**. Distribution of responses across the population relative to the mean (100%). The distributions are illustrated with half violin plots (blue and left: feedback present; red and right: feedback blocked. A “swarm-type” scatter plot is overlaid showing individual data points (46 neurons x 3 stimulus locations). The median is shown as a diamond marker, and asymmetric standard deviation bars are shown extending from the 100% mean level. Numbered stars indicate significant differences in the variance of the normalized population responses as tested with a Brown-Forsythe test, with exact values as follows: *1, p=0.009; *2, p=0.005; *3, p=0.007; *4, p=0.008. **C**. Mean firing rates of neurons that contribute most to spatial discrimination for a given stimulus pair, separated based on which stimulus location led to the strongest response. The firing rate in response to the location that drives them most strongly is not different (Wilcoxon test, p=0.102) between feedback blocked and present conditions (red and blue distributions). The firing rate in responses to the weaker stimulus locations (cyan and orange distributions) is significantly decreased (Wilcoxon test, *1, p<10^−20^). Arrows show the distributions’ medians.

The key aspect of the population responses that encodes stimulus location is the difference in response between neurons representing different receptive fields. For example, a conspecific located in front of the focal fish will cause a stronger stimulation for neurons with receptive fields near the head than near the tail. Stronger localization cues therefore mean more pronounced differences between the neurons representing weakly vs strongly stimulated areas. We examine in Figure 9B the distribution of responses across the population and how this distribution is affected by feedback. We find that there is a significant change in the distribution shape with a particularly clear shift upward in the weakest responses when feedback is blocked. Our results show that feedback increases spatial contrast across the population by causing relatively larger reductions in responses of more weakly activated neurons. We further confirm this finding by extracting the top 20% of neurons that are most heavily weighted in the discrimination analysis of Figure 9A using firing rate as a response measure. More heavily weighted neurons have responses that are more different between the two stimulus locations being compared. It could be, for example, a neuron whose receptive field falls within the hot spot of the electric image for one stimulus location but in a weakly activated area (the “background”) for the other location. Marking, for each neuron, the response to “stimulus position 1” as the strongest response of the two locations, and stimulus position 2 as the weaker one, we find a clear decrease in the weak responses but no significant decrease in the strong responses (Figure 9C). Our results therefore support the hypothesis that feedback performs a spatial filtering akin to background suppression and contrast enhancement to support efficient spatial coding.

## Discussion

### Adaptive spatiotemporal filtering of natural conspecific signals

The present results extend the established role of cerebellum-like feedback in the ELL from cancellation of simple low-frequency amplitude modulations to the processing of conspecific signals with a realistic range of carrier (beat frequency), temporal (contrast envelope), and spatial structures. As expected, feedback strongly reduced the beat-locked modulation of responses to 5-Hz signals but did not significantly alter the representation of individual cycles at 75 or 150 Hz. Slow contrast envelopes, however, were attenuated for all three beat frequencies. The magnitude of this attenuation depended on the recent stimulus sequence: stronger contrast during the preceding approximately 0.4 s predicted a larger subsequent feedback effect. Feedback also improved discrimination among stimulus locations, primarily because it reduced weak population responses more markedly than strong responses. Our analysis suggests that firing rate, and its difference between strong and weak responses within the population, is a more reliable code of spatial information, although this might not be the case for all types of stimuli and subsets of neurons (Milam and Marsat 2026). The importance of firing rate in coding by ELL neurons was also highlighted in previous research showing that firing rate increases underlie the coding and behavioral responses to specific types of envelope signals (Huang et al. 2018; Metzen et al. 2018). Interestingly, these studies suggest that these firing rates responses are enhanced by another feedback pathway, the direct pathway from nP.

The temporal and spatial effects described here are functionally related. During a social interaction, movement changes the overall contrast of the beat and the spatial gradient across the receptor array that provides cues to the position of the other fish (Fotowat et al. 2013; Milam et al. 2019; Ramachandra et al. 2026). Feedback adapts to these spatio-temporal variations in contrast by reducing the weaker background responses within the population. Thus, the indirect feedback pathway is better described as an adaptive spatiotemporal filter than as simple gain control to reduce low frequencies.

### Recent stimulus history adaptively shapes the strength of feedback

Our analysis shows that the instantaneous contrast of the current beat cycle alone cannot explain feedback strength. Beat cycles with comparable contrast were attenuated differently depending on whether they followed stronger or weaker stimulation. Using random envelopes, the estimated linear kernel related the feedback-dependent change in firing rate to contrast over a window extending up to roughly 400 ms before the response. The pathway is therefore entrained by the sensory inputs, and the feedback output adapts to the recent signal strength. This history dependence is compatible with several known features of the circuit. Parallel-fiber synapses exhibit short-term synaptic dynamics that can rapidly adjust the feedback strength (Lewis and Maler 2004). Descending inputs into the ELL can adjust neural tuning and response variability to “whiten” the responses to the envelope, therefore adjusting feedback’s impact depending on stimulus frequency (Huang et al. 2018, 2019). Furthermore, some cerebellar circuits can act as resonators for low-frequency signals (Gandolfi et al. 2013), and we recently hypothesized (Johnson et al. 2025) that such a resonator-based mechanism could underlie the frequency-specific cerebellar channels in this electrosensory cerebellar feedback pathway. A resonator-based mechanism would be consistent with both the frequency specificity previously demonstrated (Bol et al. 2011) and the history dependence that we characterize here.

The current experiments do not distinguish the contributions of possible mechanisms, but they place a useful constraint on circuit-level accounts: the effective feedback signal reflects an integration of recent stimulus strength with a decay extending over hundreds of milliseconds. In this operational sense, the pathway performs a predictive computation. Recent input changes the response to continuing or recurring stimulus components, as expected when feedback carries an estimate of the sensory pattern likely to persist (Bell et al. 2008; Keller and Mrsic-Flogel 2018).

### Naturalistic stimulation refines the negative-image model

The effects observed with constant-contrast beats are consistent with the classical negative-image model of electrosensory feedback. For low-frequency, spatially broad stimulation, parallel-fiber input is shaped so that its net effect opposes the phase-dependent feedforward response, reducing firing near the preferred phase of the beat and increasing it near the opposite phase without necessarily changing mean firing rate (Bastian 1986; Bastian et al. 2004). The frequency dependence observed here also agrees with previous work. The burst dynamics of superficial pyramidal cells, the range of delays represented in the parallel-fiber pathway, and burst-dependent synaptic plasticity together support phase-specific cancellation only over a restricted low-frequency range (Bol et al. 2011; Harvey-Girard et al. 2010). Individual cycles of 75- and 150-Hz beats do not drive bursting and phase-specific depression to generate a phase-specific negative image. Yet these high-frequency beats still contain slower variations in contrast, and ELL circuits can represent the AM beat and the low-frequency envelope within the same responses (Middleton et al. 2006). The carrier AM signal does not contain frequencies in the 0-1 Hz range: envelope modulations cause small changes in the frequency of the AM carrier (“sidebands” components of the beat-frequency spectrum peak). With simple envelope extraction mechanisms (Longtin et al. 2008), these contrast changes do cause modulations in the firing rate of pyramidal cells. Since deep pyramidal cells drive the feedback pathway, firing rate modulations caused by the envelope or by the beat are treated indiscriminately in the downstream feedback circuit.

The feedback pathway can influence responses to conspecific envelope signals in various ways (Hofmann and Chacron 2019), including by affecting response heterogeneity (Metzen and Chacron 2023), and this effect depends on the detailed characteristics of the signal (Ly and Marsat 2018). Our results extend these findings by showing that the effect of cerebellar feedback on envelope coding extends to the full range of conspecific signals, including those with higher frequency beats. However, the impact of feedback on individual neurons’ responses is not a simple enhancement of envelope coding within each neuron. Using realistic conspecific signals, we show here that a significant portion of the neurons have their response to the envelope signal attenuated by feedback. We argue that the benefits of feedback input on movement-related envelope coding have to be evaluated at the population level. Indeed, conspecific signals have specific spatial structures that have thus far been ignored and simplified to a uniform “global” spatial pattern. A realistic conspecific electric image is broader than a prey-like local stimulus, but it is not uniform: it contains a strong region near the sender and an extended background of weaker activation (Milam et al. 2019; Ramachandra et al. 2026). This gradient changes as the two fish move further apart. The present findings show that the negative-image mechanism remains engaged under this realistic geometry, albeit somewhat weaker, while revealing consequences on spatial coding by the population that would be obscured by spatially uniform stimulation.

### Graded background suppression enhances spatial coding

The spatial coding analysis identifies a functional consequence of the feedback attenuation that is not apparent from the response of a single neuron. With feedback intact, responses to the three stimulus locations were more readily discriminated, and a given level of performance could be reached with fewer neurons. The effect was present across the response measures examined and was particularly pronounced when mean firing rate was used as the aspect of neural responses encoding spatial information.

This improvement did not result from a uniform decrease in population activity. Rather, feedback changed the distribution of responses by producing larger relative reductions among weakly activated neurons while leaving the strongest responses comparatively unchanged. This pattern is well suited to the geometry of a conspecific electric image.

Neurons whose receptive fields overlap the “hot-spot” of the electrosensory image respond most strongly, whereas neurons representing portions of the skin more distant from the other fish receive weaker feedforward drive akin to a background activation level. The spatially broad integration of the indirect feedback pathway provides a plausible basis for suppressing this distributed background (Berman and Maler 1999; Chacron et al. 2005; Sas and Maler 1987, 1987). A feedback input reflecting the overall signal strength can have a proportionally stronger impact on weak background responses than on the responses of neurons strongly driven by feedforward inputs. As a consequence, it increases the difference between neurons representing strongly and weakly stimulated regions, thereby making population patterns associated with different source locations more distinct.

### Feedback and predictive signal shaping efficient population codes

The temporal and spatial findings in this paper can be understood as two consequences of the same circuit organization. A feedback signal based on recent, spatially distributed activity would be recruited most strongly by components that are both persistent in time and broadly represented across the population. Subtracting that estimate from current input would lead the responses to represent only changes in the signal: differences across space and time. Differences across space signal fish location, whereas differences across time signal movement of the fish. This draws a concrete link between predictive and efficient coding. Prediction refers here to the dependence of feedback on recent input characteristics. In contrast, efficient coding refers to the resulting allocation of population dynamic range away from common or redundant activity and toward variation that distinguishes stimulus conditions. The present data do not directly quantify information maximization or prediction error, but they show that an identified descending pathway can use temporal context to reshape the distribution of activity across a sensory population. This circuit-level operation is consistent with both adaptive redundancy reduction in the ELL (Bastian et al. 2004) and broader formulations linking efficient and predictive neural coding (Manookin and Rieke 2023; Pitkow and Meister 2012).

Natural signals rarely vary along a single dimension. They contain local and global structure, fast fluctuations nested within slower envelopes, and correlations that extend into the recent past. Feedback that uniformly suppresses or amplifies an entire sensory representation would be poorly suited to this structure. The present results instead illustrate how adaptive suppression can improve signal representation by reducing components that are shared across space or time while retaining differences that are more useful for discrimination. Comparable computations are common across sensory systems, even when the underlying anatomy differs. In the retina, wide-field inhibitory circuits reduce responses to motion shared by an object and its background, allowing relative object motion to remain prominent (Baccus et al. 2008). In auditory pathways, contralateral inhibition enhances differences between bilateral channels, and its effect can depend on whether the temporal structure of excitation and inhibition is matched to the natural signal (Marsat and Pollack 2005). Predictive influences in cortical sensory pathways similarly alter responses according to context and prior experience rather than acting as a fixed change in gain (Keller and Mrsic-Flogel 2018). The electrosensory system adds an important mechanistic example and shows the subtle and complex shaping that feedback imposes on the sensory stream. It improves the organization of activity, attenuating predictable and broadly shared components so that the remaining population response more clearly represents distinctions among natural stimuli.

## Acknowledgements, & Statements

We thank Drs Ramachandra and Milam for training and guidance on experimental procedures.

## Author Contributions

GM conceived and designed the research. DM, ES and BH performed experiments. DM, ES, and GM analyzed data, interpreted results of experiments, prepared figures, drafted the manuscript, edited and revised the manuscript, and approved the final version of the manuscript.

